# The impact of Hurricane Michael on longleaf pine habitats in Florida

**DOI:** 10.1101/736629

**Authors:** Nicole E. Zampieri, Stephanie Pau, Daniel K. Okamoto

## Abstract

The longleaf pine (*Pinus palustris*) ecosystem of the North American Coastal Plain (NACP) is a global biodiversity hotspot. Disturbances such as tropical storms play an integral role in ecosystem maintenance in these systems. However, altered disturbance regimes as a result of climate change may be outside the historical threshold of tolerance. Hurricane Michael impacted the Florida panhandle as a Category 5 storm on October 10th, 2018. In this study, we estimate the extent of Florida longleaf habitat that was directly impacted by Hurricane Michael. We then quantify the impact of Hurricane Michael on tree density and size structure using a Before-After study design at four sites (two wet flatwood and two upland pine communities). Finally, we identify the most common type of tree damage at each site and community type. We found that 39% of the total remaining extent of longleaf pine habitat was affected by the storm in Florida alone. Tree mortality ranged from 1.3% at the site furthest from the storm center to 88.7% at the site closest. Most of this mortality was in mature sized trees (92% mortality), upon which much of the biodiversity in this habitat depends. As the frequency and intensity of extreme events increases, management plans that mitigate for climate change impacts need to account for large-scale stochastic mortality events in order to effectively preserve critical habitats.

## 1. Introduction

Disturbance plays an integral role in maintaining ecosystem structure and functioning (1,2). However, many ecological disturbances are expected to change as the climate changes (3), altering the frequency, intensity, duration, and timing of events (4). Shifting disturbance regimes due to climate change pose a threat to the conservation of biodiversity as species experience conditions outside their historical norms (5–7). In forest and savanna ecosystems, disturbances can include fire, hurricanes, extreme wind events, insect outbreaks, exotic plant invasions, or drought, among many others (1).

Longleaf pine (*Pinus palustris* Mill.) forests and savannas provide critical habitat for numerous endangered species of animals such as the red-cockaded woodpecker (*Picoides borealis*), gopher tortoise (*Gopherus polyphemus*), and the indigo snake (*Drymarchon corais couperi*) as well as many endangered plants such as the American chaffseed (*Schwalbea americana*), Florida skullcap (*Scutellaria floridana*), and Harper’s beauty (*Harperocallis flava*) (8–11). About 30% of all plant species associated with longleaf pine habitat are endemic to the region (12). Yet, the range of longleaf pine has been reduced to <3% of its historical extent (13). Florida, and more specifically the Florida Panhandle, is one of the most important strongholds of endangered longleaf pine habitat (14,15) containing 51% and 31%, respectively, of all the remaining longleaf pine ecosystem (16,17).

Longleaf pine habitats in the Panhandle of Florida are located within the North American Coastal Plain (NACP) global biodiversity hotspot (12). The NACP borders the Gulf and Atlantic coast, which is subject to frequent storm events (12). Over the course of a century, the entire range of the NACP will have experienced at least one major hurricane (Category 3 and above) (18–20). There have been numerous studies assessing damage to forests in the NACP after major storm events (e.g., Gresham et al. 1991; Xi et al. 2008; Johnsen et al. 2009; Kush and Gilbert 2010; Dyson and Brockway 2015). Longleaf pine trees have been found to have lower mortality than other species when exposed to hurricane force winds (23,24,26–28). Species that evolved within the coastal plain have been shown to have lower mortality than species whose evolutionary range extends beyond the coastal plain region, possibly due to strong selection pressure from frequent exposure to high wind storms over their lifetime (23,27). However, as the climate changes, high wind storm events such as hurricanes and tornadoes will increase in strength and/or frequency, outside of the system’s historic norms (4,29–31).

Management of longleaf pine ecosystems is generally aimed at conserving and expanding the extent of mature, open-canopied habitat maintained by frequent fire (17,32). The highest quality stands are considered to be mature forests with a frequent enough fire regime to promote regeneration of longleaf and maintain a highly biodiverse understory – estimated at <0.5% of its historical coverage (13). These systems have ranging tree basal areas between <100 to 300+ trees·ha^−1^ (33) and require frequent fire (1-5 year return interval) (11,13,34,35). Canopy gaps promote a biodiverse understory (36) and allow for recruitment and regeneration of longleaf pine (37). These gaps in the canopy produced by fallen trees allow for greater light penetration and colonization by shade-intolerant species (19,38). Most successful recruitment of longleaf pine requires patches in the canopy to be opened up by disturbances such as fire, wind, or rain events (37,39).

Hurricanes may contribute to necessary gap dynamics by removing older, rotten trees and other species that may be crowding out the understory (1,19,27,39–41). While gap dynamics driven by storm events play an important role in maintaining these open-canopied habitats, the potential for hurricanes of increasing strength to occur over the next century (4,29,30) combined with the lack of remaining habitat (12,14) could lead to severe damage and potentially permanent losses of remnant stands of an already vulnerable system. The resilience of each stand will depend on localized conditions including the availability of a seed source and active habitat management that allows establishment and survival of longleafs (7). The loss of mature trees and severe damage to the understory may impede natural regeneration, alter the fire regime, increase the chance of invasive species establishment, and provide favorable conditions for insect outbreaks (3,4,42–45).

On October 10^th^, 2018 Hurricane Michael made landfall in the Florida Panhandle as the first Category 5 storm on record in the region. It was the strongest hurricane to make landfall in the continental U.S. since Hurricane Andrew in 1992 with maximum sustained winds of 257 km/h and minimum barometric pressure of 919 mb (Beven II et al., 2019, National Hurricane Center). Here we investigate the impact of Hurricane Michael on four longleaf pine habitats in the Florida Panhandle through a Before-After assessment (46) of tree density and size structure. We first determine the extent of longleaf pine habitat in Florida affected by Hurricane Michael. We then classify and compare longleaf damage (e.g. uprooted, snapped, crown damage) and mortality at each site, and discuss implications for management and restoration.

## 2. Methods

### 2.1 Hurricane Coverage and Extent of Impacted Habitat

Data on the storm track and wind extent was obtained from the National Hurricane Center. Hurricane force winds extended outward from the storm center for 75 km and tropical storm force winds extended 280 km (47).

Using ArcMap 10.6.1, we created buffers around the storm track for hurricane and tropical storm force winds. We then overlaid the buffers on longleaf pine habitat coverage within Florida obtained from the Longleaf Pine Ecosystem Geodatabase (LPEGDB) (https://www.fnai.org/longleafgdb.cfm). The LPEGDB is a publicly available geodatabase with extensive data on the distribution and ecological condition of longleaf pine habitat in Florida. Pinelands were identified using aerial images, data provided by agencies, field surveys, and parcel data. Pinelands were then classified by longleaf pine occurrence as “known”, “expected”, “potential”, or “pinelands other than longleaf”. “Known” habitat has been confirmed through field surveys, “expected” are expected to be longleaf dominated based on historical vouchers, natural community type, and/or presence of red-cockaded woodpeckers, and “potential” are identified as having a community type that may be suitable for longleaf but there are no records of presence and further assessment is needed (16). We then extracted the area of known, expected, and potential longleaf habitat within the hurricane force and tropical storm force wind buffers to determine the extent of habitat impacted by the storm within Florida.

### 2.2 Site Description

In the summer of 2018, pre-Hurricane Michael, we surveyed several ‘exemplary’ longleaf pine reference sites (48) throughout the state of Florida to assess longleaf pine density, age and size structure. Four of these initially surveyed sites were in the path of Hurricane Michael and are the focus of the Before-After assessment in this study. The Florida Natural Areas Inventory (FNAI) selected individual sites to serve as reference sites based on canopy structure, regeneration, and overall groundcover quality, relative to pre-Columbian conditions. The longleaf pine community reference sites are well managed (with active fire management), exemplary representations of their respective community types and are mostly comprised of second-growth stands of naturally occurring longleaf pine (16,48). The four sites in this study represent two different natural community types, wet flatwoods (WF) and upland pine (UP), ranging between 2 and 85 km away from the center of the storm (Fig 1). The two WF sites were in St. Marks National Wildlife Refuge (NWR) (85 km from center of storm) and Apalachicola National Forest (NF) (35 km from center of storm). The two UP sites were in Joe Budd Wildlife Management Area (WMA) (56 km from center of storm) and Apalachee Wildlife Management Area (WMA) (2 km from center of storm). Apalachee WMA (SUO-57197, Florida Fish and Wildlife Conservation Commission), Joe Budd WMA (SUO-57198, Florida Fish and Wildlife Conservation Commission), and St. Marks NWR (SUP FF04RFSM00-2018-0013, U.S. Fish and Wildlife Service), granted permitted access to perform field research. No permit was required for access to the Apalachicola NF site (Kelly Russell, Forest Supervisor, National Forests in Florida, United States Department of Agriculture).

**Fig 1.**
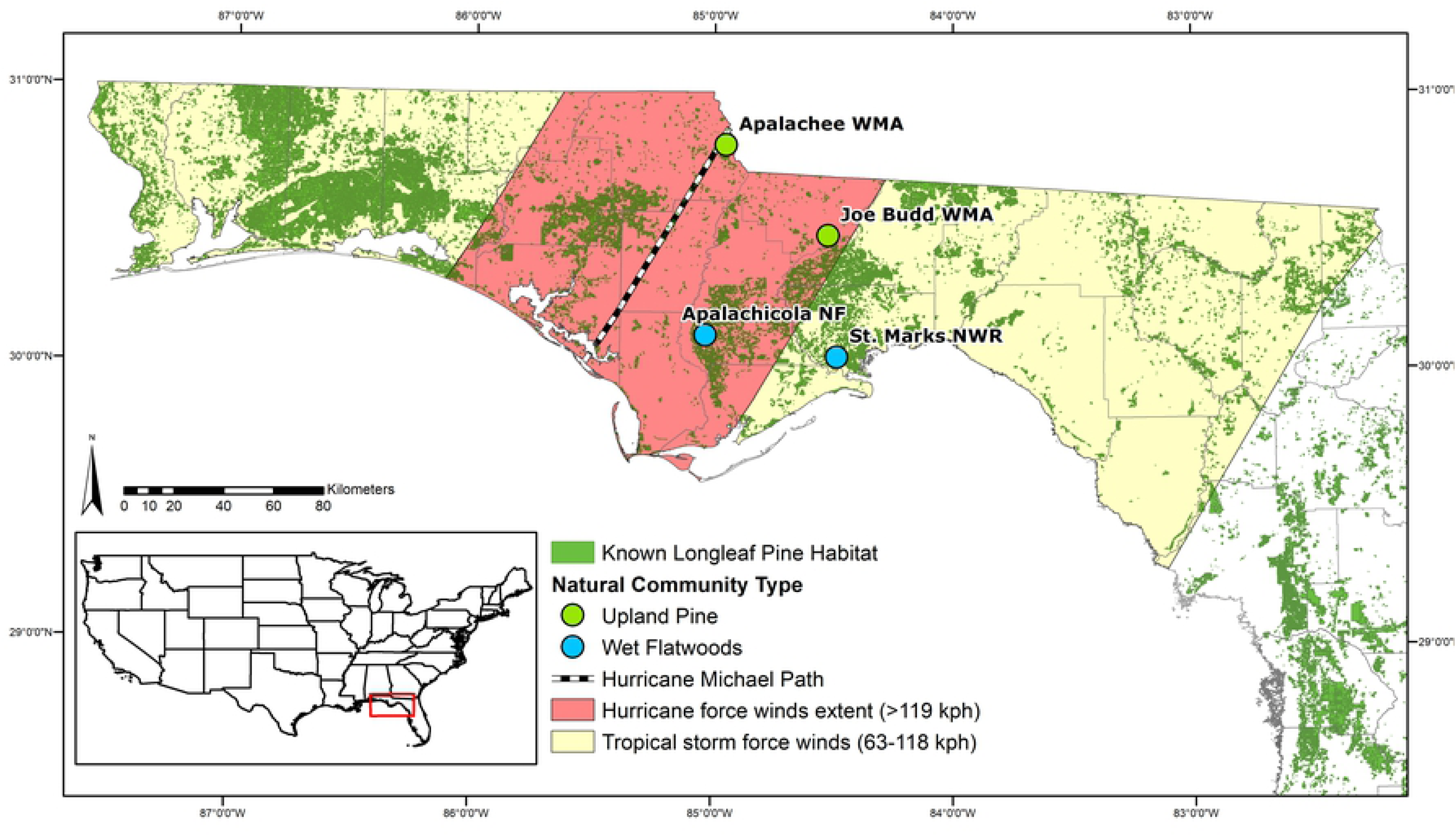
Map of study sites and storm coverage. The four study sites in the Florida Panhandle in the path of Hurricane Michael include: Apalachee WMA, Joe Budd WMA, Apalachicola NF, and St. Marks NWR. The “known” longleaf pine habitat is extracted from the LPEGDB (Florida Forest Service and Florida Natural Areas Inventory, 2018)

Longleaf pine stands are generally monotypic, with no other species making up the dominant canopy. The systems are largely open canopy, with an herbaceous, grass dominated understory (40,49,50). Frequent seasonal fire is an integral part of this ecosystem and may be the most important process in maintaining ecosystem structure and assemblage (12,51–54). Longleaf pines have a unique life history with a “grass stage” where saplings do not put on any vertical growth for anywhere between 1-20 years (55), one of their many adaptations to fire (56). The wet flatwoods sites are more savanna-like than the upland pine sites with a very open canopy and widely spaced pine trees. The upland pine sites have trees that are closer together and include a midstory of infrequent oaks (*Quercus* spp.). In contrast to WF sites, UP sites are dry, well drained, and have a greater distance between the water table and the surface (49). These differences in soils and hydrology may affect their response to high wind events (57,58).

### 2.3 Pre- and Post-Hurricane Field Surveys

Prior to the hurricane, sites were surveyed in April and May of 2018. Field surveys of tree density, life-stage, and size structure were conducted using modified variable area transects (59). A baseline transect was extended 40 meters and divided into 8 cells (4 on each side, each 10 m wide and variable in length) to make a plot. Within each cell, data on the closest 5 living trees were recorded, including GPS location, diameter at breast height (dbh), and distance to the furthest tree, for a maximum of 5 trees per cell or a maximum search distance of 20 m per cell. The number of plots varied from 2-5 depending on the size of the stand, to capture a representative sample of each site. Trees were classified into 5 possible size classes based on their life stage and dbh: grass stage, juveniles (<15 cm dbh), younger mature (15-30 cm dbh), mature (30-45 cm dbh), or older mature (45+ cm dbh).

Post-hurricane surveys were conducted in November and December of 2018, within 3 months of the storm, using the same variable area transect methodology. Plots were relocated using GPS. Although transect placement matching prior surveys was not exact, the variable-area transects are designed to capture representative density estimates for the site. During post-hurricane surveys, additional information was recorded, including the status of the tree (living or dead) and any visible damage. Post-hurricane surveys were conducted two ways. First, a survey of remaining living trees was conducted for the Before-After assessment of tree density. Second, a survey of all trees (living and dead) was conducted to determine the density of dead trees as well as percent mortality. Living and dead trees were classified into the following damage groups: no visible damage, minor damage (such as needle loss, broken, or fallen branches), partially uprooted, uprooted, snapped, or moderate to major crown damage, for which percent canopy loss was also recorded (canopy loss of >50%, >75%, or >90%). Trees that were partially uprooted, uprooted, snapped, or had canopy loss of 75% or greater were considered dead for our mortality assessment. Canopy loss of 75% or greater included damage to the main stem and majority needle loss. Canopy loss of 90% included damage to the main stem and total needle loss. Only trees that died as a result of the storm were included in the survey. Those that looked diseased prior to the storm or had signs of decay inconsistent with other trees were not included. Grass stage individuals were classified as living or dead but were excluded from the damage classification. Grass stage individuals were considered dead when there was visible death to the apical meristem (usually crushed and/or black-brown).

We quantified the effect of the hurricane on tree density in two ways. First, we compared densities in pre- and post-hurricane surveys, and second, we directly estimated mortality by comparing the density of living and dead trees post-hurricane. For the former, we estimated densities of pre- and post-hurricane trees by size class using generalized linear mixed effects models, where site and the interaction between site and survey (i.e., before vs. after) were fixed parameters and sample plot within site was a random effect. To estimate mean longleaf pine mortality at each site, we used a generalized linear mixed-effects model allowing mortality estimates to vary randomly among sample cells within plots. In the density estimates, plots were used as the random effect because not every size class was represented in every cell, whereas in the mortality estimates, mortality was aggregated across size classes, and cells within plots were the random effect. We also report per capita mortality observations by size class at each site (determined as number of observed dead trees over the total number of trees per size class). The grass stage was excluded from mortality estimates because their deaths could not be directly attributed to the hurricane.

## 3. Results

### 3.1 Hurricane Coverage and Extent of Impacted Habitat

Within the Florida Panhandle, the storm impacted between 533,000 to 1,043,000 hectares of longleaf pine habitat. Tropical storm force winds impacted a total of 533,000 “known” longleaf pine habitat. An additional 15,000 ha of “expected” longleaf and 495,000 ha of “potential” longleaf were within the tropical storm force winds (280 km buffer). Hurricane force winds (75 km buffer) impacted 114,000 ha of “known” longleaf pine habitat. An additional 4,000 ha of “expected” longleaf and an additional 54,000 ha of “potential” longleaf were within the hurricane force winds buffer.

### 3.2 Wet Flatwoods (WF)

#### 3.2.1 St. Marks National Wildlife Refuge

St. Marks NWR, the site furthest from the storm center (85 km), had the least amount of damage recorded. This site had the highest density of grass stage individuals, 236 (SE = 85) and 234 (SE = 49) trees·ha^−1^ pre- and post-hurricane respectively (Table 1). Mature trees were only represented by the younger mature size class (15-30 cm dbh). Overall tree density (including grass and juvenile stage) decreased by 0.6% from 331 (SE = 76) to 329 (SE = 40) trees·ha^−1^. Mature tree density did not show a significant decrease (from 71 (SE = 17) to 81 (SE = 18)). Only the juvenile size class showed a significant decrease (from 24 (SE = 9) to 14 (SE = 9)) (Table 1). The overall mean tree densities were similar pre- and post-hurricane (Fig 2). This site had the lowest estimated mortality of 1.3% (95% CI: 0.12 – 5.6%) (Table 2). All trees that died were snapped (Table 2) and in the younger mature size class (Fig 3).

**Table 1.** Density assessment of longleaf pine trees Before-After Hurricane Michael.

|  | Pre-hurricane |  |  |  | Post-hurricane |  |  |  |
| --- | --- | --- | --- | --- | --- | --- | --- | --- |
|  | St. Marks<br>NWR (WF) | Apalachicola<br>NF (WF) | Apalachee<br>WMA (UP) | Joe Budd<br>WMA (UP) | St. Marks<br>NWR (WF) | Apalachicola<br>NF (WF) | Apalachee<br>WMA (UP) | Joe Budd<br>WMA (UP) |
| Grass Stage | 236 (85) | 42 (13) | 63 (32) | 2 (2) | 234 (49) | 45 (18) | **10 (7)** | 2 (2) |
| Juveniles (<15 cm dbh) | 24 (9) | 17 (9) | 19 (6) | 19 (7) | **14 (9)** | 18 (10) | **5 (2)** | **17 (5)** |
| Younger Mature (15-30 cm dbh) | 71 (17) | 26 (10) | 14 (5) | 71 (14) | 81 (18) | 37 (10) | 2 (2) | 82 (13) |
| Mature (30-45 cm dbh) | 0 | 22 (7) | 62 (8) | 144 (19) | 0 | 12 (6) | **2 (2)** | 102 (16) |
| Older Mature (45+ cm dbh) | 0 | 0 | 26 (5) | 11 (4) | 0 | 0 | **3 (2)** | 21 (8) |
| Overall Mature Tree Density | 71 (17) | 48 (9) | 102 (10) | 225 (23) | 81 (18) | 50 (10) | **8 (4)** | 206 (21) |
| Overall Living Tree Density | 331 (76) | 108 (25) | 184 (30) | 246 (24) | 329 (40) | 113 (18) | **23 (9)** | 224 (21) |
| Dead Tree Density |  |  |  |  | 4 (4) | 9 (0) | 128 (5) | 7 (3) |
| Percent Change in Mature Tree Density |  |  |  |  | 14.1% | 4.2% | -92.2% | -8.4% |
| Percent Change in Overall Density |  |  |  |  | -0.6% | 4.6% | -87.5% | -8.9% |

Values are reported in trees·ha<sup>-1</sup> with standard error in parentheses. Post-hurricane densities with a significant decrease from pre-hurricane densities per size class and overall at p-value < 0.01 are bolded. Pre-hurricane surveys only included living trees. Percent change in mature tree density includes the younger mature, mature, and older mature size classes. Percent change in tree densities are different from our mortality estimates (Table 2) because mortality estimates were obtained using a mixed-effects model that weights density data from each cell in the

**Table 2.** Damage Classification and Mortality.

|  | St. Marks NWR<br>(WF) | Apalachicola NF<br>(WF) | Apalachee WMA<br>(UP) | Joe Budd WMA<br>(UP) |
| --- | --- | --- | --- | --- |
| No Visible Damage | 56 (93.3%) | 29 (51.8%) | 12 (8.3%) | 128 (82.6%) |
| Minor | 0 | 18 (32.1%) | 4 (2.8%) | 19 (12.3%) |
| Partially Uprooted | 0 | 2 (3.6%) | 6 (4.1%) | 0 |
| Uprooted | 0 | 7 (12.5%) | 46 (31.7%) | 0 |
| Snapped | 4 (6.7%) | 0 | 70 (48.3%) | 7 (4.5%) |
| Canopy Loss >50% | 0 | 0 | 1 (0.7%) | 1 (0.6%) |
| Canopy Loss >75% | 0 | 0 | 4 (2.8%) | 0 |
| Canopy Loss >90% | 0 | 0 | 2 (1.4%) | 0 |
| Estimated Mortality | 1.3% (0.12 - 5.6%) | 8.4% (1.8 - 23.2%) | 88.7% (78.8 - 96.0%) | 3.1% (0.7 - 8.1%) |

**Fig 2.**
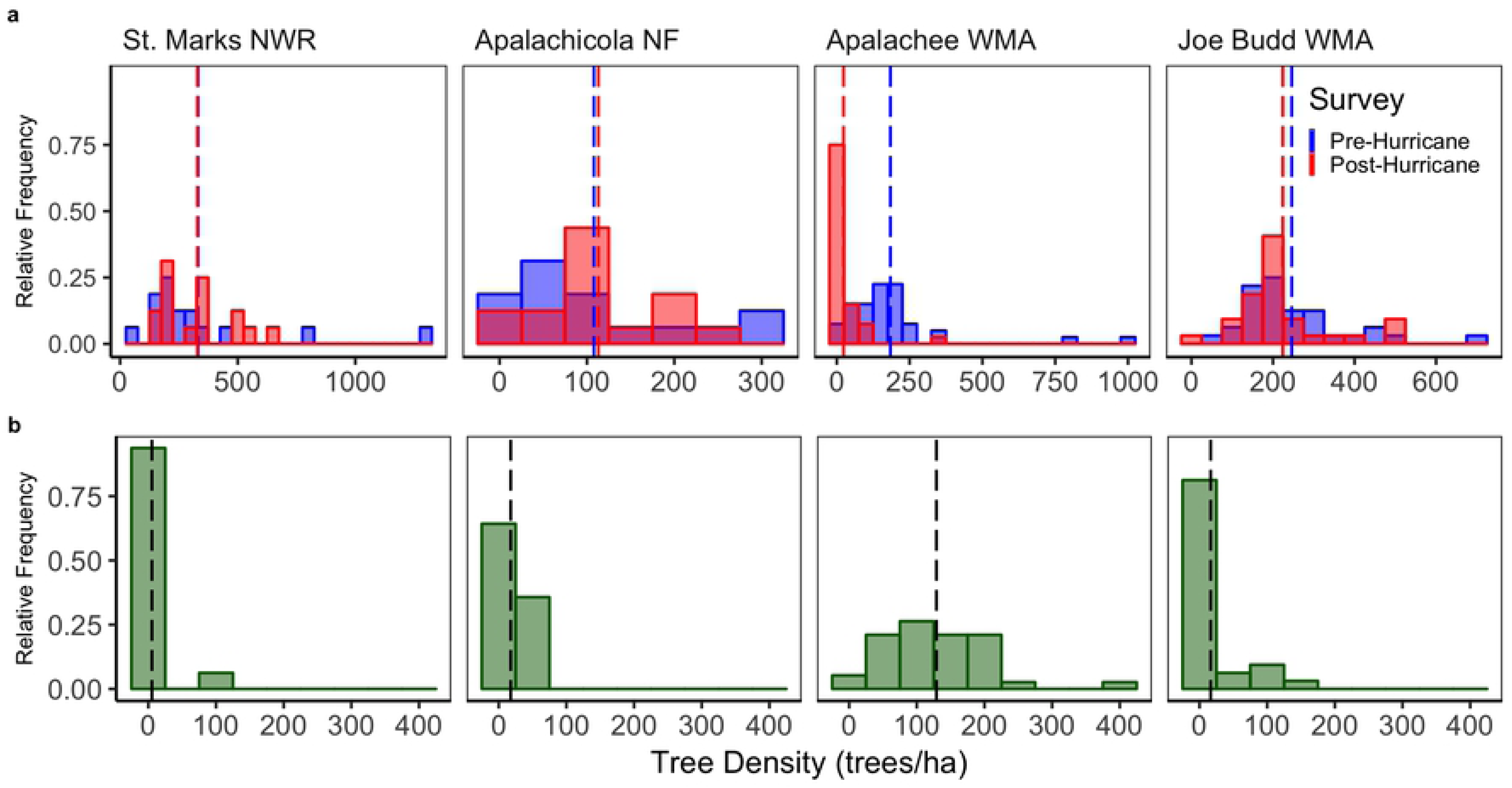
Pre- and Post-Hurricane Tree Density and Observed Tree Mortality. a. Histograms of pre- and post-hurricane living tree densities from each cell in all transects show the most dramatic change in tree density at Apalachee WMA, whereas other sites show less change or no detectable change. Group means of living tree density are indicated by dashed lines. Each site is scaled on a different x-axis for clearer visualization. b. Histograms of observed tree mortality show densities of dead trees from each cell in all transects at all sites post-hurricane. The mean overall dead tree densities are indicated by dashed lines

**Fig 3.**
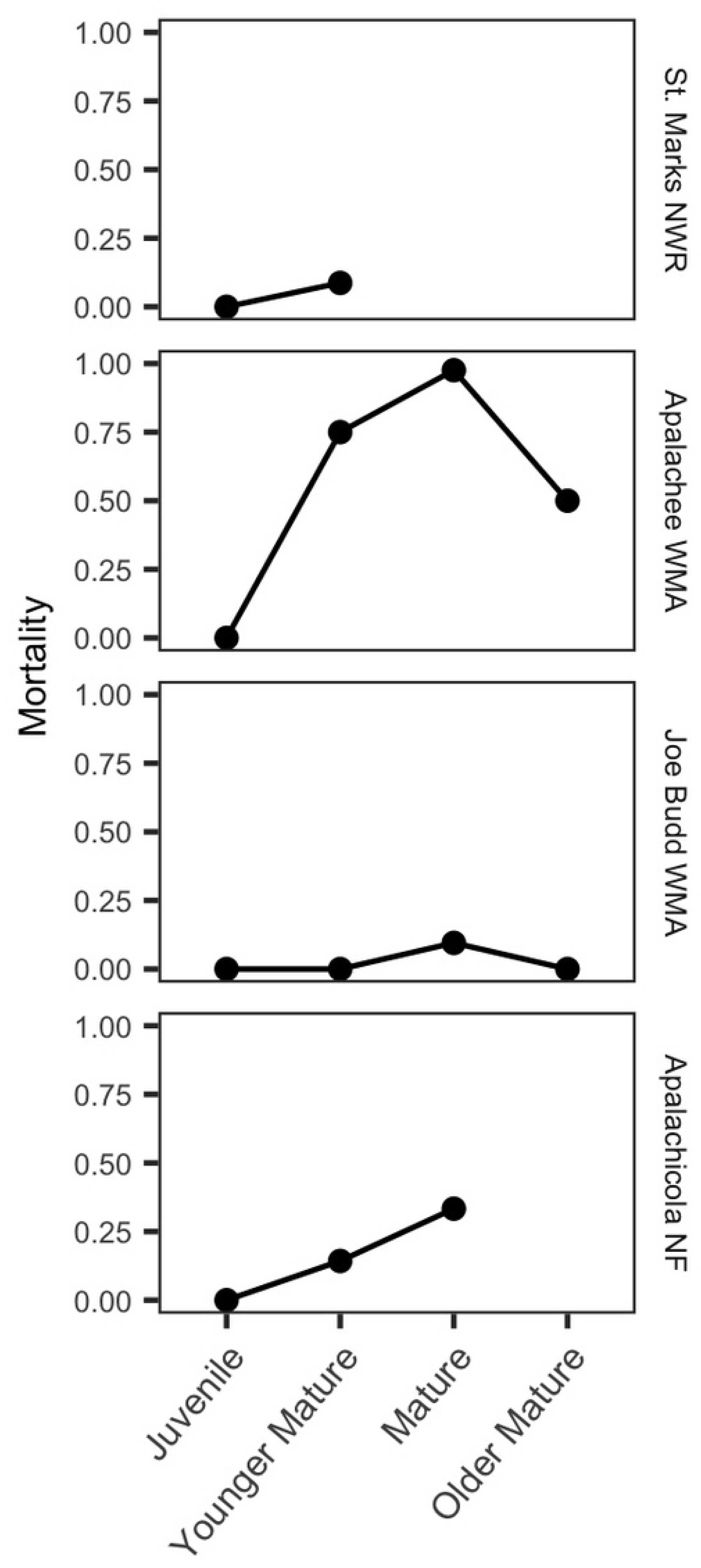
Percent mortality relative to overall mortality within each size class of longleaf pine at four sites. Size classes are as follows: juveniles (<15 cm dbh), younger mature (15-30 cm dbh), mature (30-45 cm dbh), or older mature (45+ cm dbh)

Values are reported in trees·ha^−1^ with standard error in parentheses. Post-hurricane densities with a significant decrease from pre-hurricane densities per size class and overall at p-value < 0.01 are bolded. Pre-hurricane surveys only included living trees. Percent change in mature tree density includes the younger mature, mature, and older mature size classes. Percent change in tree densities are different from our mortality estimates (Table 2) because mortality estimates were obtained using a mixed-effects model that weights density data from each cell in the variable-are transects for a site level mean.

The damage classification included both living and dead trees and did not include grass stage individuals. Values are reported in trees·ha^−1^ followed by the total percentage from each site. Trees were classified as follows: no visible damage, minor damage (minor visible damage such as needle loss or fallen branches), partially uprooted, uprooted, snapped, or minor to major crown damage including canopy loss of >50%, >75%, or >90%. Estimated site level mortalities included all size classes and were determined in the generalized linear mixed effects model. 95% confidence intervals are presented in parentheses. Trees that were partially uprooted, uprooted, snapped, or had canopy loss of 75% or more are included in the total estimated mortality.

#### 3.2.2 Apalachicola National Forest

Apalachicola NF was closer to the storm center than St. Marks NWR at 35 km away. This site experienced slightly higher mortality and had a higher density of damaged trees than St. Marks NWR (Table 2). The site had trees in all size classes except the largest size class (45+ cm dbh). Overall tree density (including grass and juvenile stage) increased by 4.6% from 108 (SE = 25) to 113 (SE = 18) trees·ha^−1^. Overall and mature tree density at Apalachicola NF did not show significant decreases (from 102 (SE = 25) to 113 (SE = 18) trees·ha^−1^ and from 48 (SE = 9.0) to 50 (SE = 10) trees·ha^−1^, respectively) (Table 1). Estimated mortality was 8.4% (95% CI: 1.8 – 23.2%). All trees that died were uprooted or partially uprooted (Table 2). Mortality across size classes shows greater mortality in larger size classes; up to 14% in the younger mature size class and 33% in the mature size class (Fig 3).

### 3.3 Upland Pine (UP)

#### 3.3.1 Joe Budd Wildlife Management Area

Joe Budd WMA is situated 56 km away from the storm center. At this site, trees were found in all size classes, including older mature trees. This site experienced greater overall loss in tree density than the WF sites but had less mortality than Apalachicola NF (WF). Overall tree density (including grass and juvenile stage) decreased by 8.9% from 246 (SE = 24) to 224 (SE = 21) trees·ha^−1^. Mature tree density decreased by 8.4% from 225 (SE = 23) to 206 (SE = 21) trees·ha^−1^ (Table 1). Only the juvenile size class showed a significant decrease (from 19 (SE = 7) to 17 (SE = 5)). All trees that died were snapped and there was 3.1% (95% CI: 0.7 – 8.1%) mortality (Table 2). Across size classes, relative mortality was higher in the mature size class (10%) than in other size classes (0%) (Fig 3).

#### 3.3.2 Apalachee Wildlife Management Area

Apalachee WMA is located 2 km away from the center of the storm and was the most severely impacted by the storm (see Figure 4). All size classes were represented at the site, including older mature trees. Overall tree density (including grass and juvenile stage) decreased by 87.5% from 184 (SE = 30) to 23 (SE = 9) trees·ha^−1^. Mature tree density decreased by 92.2% from 102 (SE = 10) to 8 (SE = 4) trees·ha^1^. Grass stage individuals were also severely impacted, entirely missing from most cells. The density of grass stage individuals decreased from 63 (SE = 32) to 10 (SE = 7) trees·ha^−1^ (Table 1). All size classes had a significant decrease in density (p-value <0.01) except the younger mature class. Total estimated mortality at the site was 88.7% (95% CI: 78.8 – 96.0%). Almost all trees at this site had some amount of visible damage and tree death was most commonly by snapping (48.3%) (Table 2). However, surviving longleafs were almost entirely grass stage and juvenile trees, and mortality increased towards the mature size class (98%), which then dropped at the older mature size class (50%; Fig 3).

**Fig 4.**
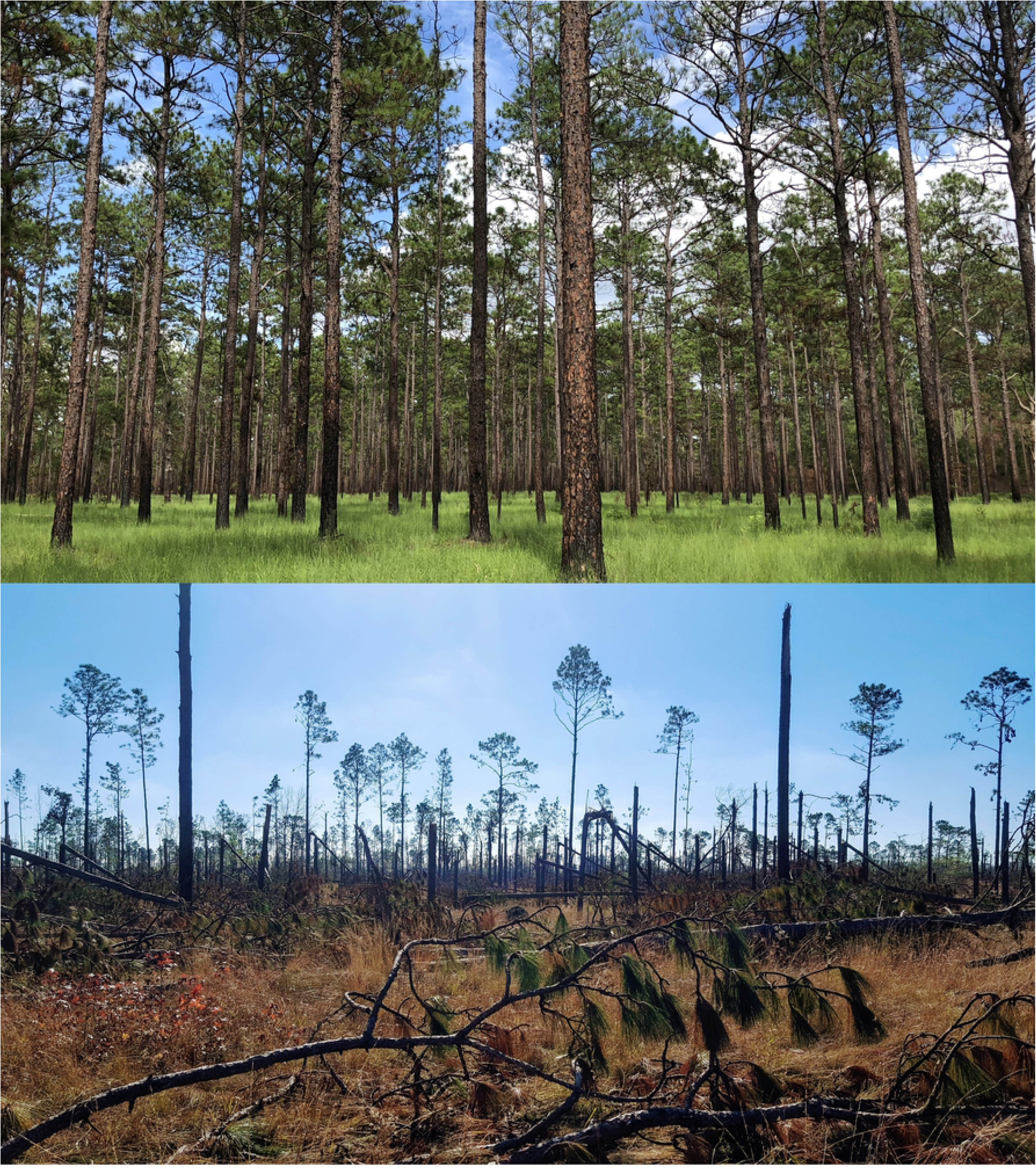
Apalachee WMA. Pre-hurricane, July 7^th^, 2018 (top, image: C. Anderson) and post-hurricane, December 1^st^, 2018 (bottom, image: N. Zampieri)

## 4. Discussion

### 4.1 Extent of Hurricane Michael’s impact in Florida

The Florida Panhandle is a stronghold for the longleaf pine system, with more connected, protected longleaf pine habitat than anywhere else in its range (16,17). Considering that the total range of longleaf pine habitat is 1.4 million ha (17), our results show that 39% and 8% of all remaining longleaf pine habitat experienced tropical storm and hurricane force winds, respectively, in Florida alone. Given the estimates that also include expected and potential longleaf habitat, the total extent impacted by at least tropical storm force winds could be up to 1,043,000 ha (76% of all remaining habitat), and up to 172,000 ha (13% of all remaining habitat) impacted by hurricane force winds. These estimates provide a baseline to assess longleaf pine conditions because varying degrees of habitat integrity and vulnerability to storm damage and climate change exist within this range. Understanding the extent of habitat impacted by one storm event highlights the importance of conserving habitat over a broad range since unexpected losses could be high in areas affected by extreme events such as Hurricane Michael.

### 4.2 Density and mortality of longleaf post-Hurricane Michael

Our surveys show a gradient of little to severe damage of longleaf pine habitats due to Hurricane Michael depending on their distance from the storm center (Fig 1 and 2). Apalachee WMA, an upland pine site closest to the path of Hurricane Michael, experienced longleaf mortality of 88.7%, predominantly in mature size classes (Figures 3 and 4). Mature trees had 98% mortality, similar to other catastrophic hurricanes. After Hurricane Hugo (Category 4, 1989), second-growth stands of longleaf in South Carolina had 95% adult tree mortality (60). Hurricane Kate (Category 3, 1985) resulted in over 20% mortality of adult longleaf from an old-growth stand, with effects continuing for at least 5 years post-hurricane (39). The significant loss of mature trees reduces the current extent of mature habitat, on which many critically endangered species depend (11). While the remaining juveniles could represent the potential for recovery, this depends on substantial efforts to remove fallen trees and debris, managing potential pests and invasive species establishment, in addition to maintaining fire (see Section 4.3). Even then, recovery could take decades for juveniles to reach mature size classes (Figure 4). At St. Marks NWR, Joe Budd WMA, and Apalachicola NF, tree loss was much lower (1.3, 3.1, and 8.4% mortality, respectively) and similar to background mortality rates driven by lightning (39,50,61). These lower rates of mortality are reasonable for maintaining open canopy gap dynamics (7,37,39). At St. Marks NWR and Apalachicola NF, the standard errors around mean tree densities were high. Thus, the apparent increase in densities is due to high variability across cells at these sites because mortality was identified at all sites.

Although mortality was not nearly as high at St. Marks NWR, Joe Budd WMA, or Apalachicola NF sites compared to Apalachee WMA, snapped, uprooted, and minor damage were still apparent at these sites. The natural communities examined in this study – wet flatwoods and upland pine – differ in community structure, hydrology, and soil type (49) that likely play a role in the type of tree damage caused by high wind events. WF sites have a hydroperiod which causes them to be inundated for parts of the year, and the water table is relatively close to the surface (49). Trees in these systems may develop a shorter taproot and are therefore less stable (62). These trees may be more likely to be uprooted in high wind events. In contrast, upland pine sites are dry, well drained, and have a greater distance between the water table and the surface. Trees in these systems develop deeper taproots in order to reach the water table, which may also provide greater structural support in high wind events (39,49,62). These trees are more likely to snap or have damage to the crown than to uproot. Trees that are uprooted cause soil disturbance that may facilitate the establishment of invasive nonnative species (1,44,45). These trees also remain greener for longer than snapped trees since their roots may still be in contact with the water table (1,4,42,43). Snapped trees create less soil disturbance but increase the amount of dead biomass on the ground that dries more rapidly than an uprooted tree, which can create hazardous fire conditions (11). Due to the differing community characteristics, we expected more trees in WF sites to be uprooted than to experience snapping or crown damage. Our damage classification (not including grass stage individuals) generally corroborated our expectations, except in the case of St. Marks NWR (WF). This site was the furthest from the storm center (85 km) and experienced low mortality overall (1.3%). In our sample, dead trees were snapped. At the other WF site (Apalachicola NF), all dead trees were uprooted or partially uprooted. At the UP sites, as expected, trees were more likely to be snapped than uprooted. All the trees that died at Joe Budd WMA (56 km away) were snapped and at Apalachee WMA the most common cause of mortality was snapping (55%), followed by uprooting (36%).

The relationship between size classes and mortality showed that in general, mortality increased towards the mature size class, and then decreased in the older mature size class when those size classes were present. Older mature trees were the least represented in the study, only found at the UP sites. At Apalachee WMA and at Joe Budd WMA, mortality was highest in mature individuals and decreased in the older mature class by 50-100% (Fig 3). The surviving individuals in the older mature class could have traits that have enabled their survival thus far and therefore are more resilient to high winds (e.g. a deeper taproot, less lower branches contributing to structural imbalance, or differences in wood density) (41,57,63). Mortality was always higher in the mature size class than in the juvenile and younger mature size-classes. In a study of hurricane-induced mortality of longleaf pines in South Carolina, similar results were found with lower mortality (<20%) in juvenile-younger mature size classes than in mature size classes, which had up to 95% mortality. At an old-growth stand in Georgia and at a stand of south Florida slash pine (*Pinus elliottii* var. *densa*) hurricane induced mortality was also higher in the larger size classes (39,64). High mortality in the mature size class can affect regeneration potential after disturbances – fewer mature trees of reproductive age means fewer opportunities for recruitment.

Since longleaf pines are the dominant and often the only canopy species in these systems, their mortality is important for creating gaps in the canopy (37,39). Currently, lightning is considered to be the primary cause of mortality in longleaf pines and therefore is seen as the main driver of gap dynamics (61,65,66). However, Platt and Rathbun (1992) found that the rate of mortality due to hurricanes exceeded that of lightning strikes when considering a longer timeframe (e.g., 10 years) at an old-growth site. In another study in Florida, lightning mortality of longleaf pine was found to be 2.94 trees·ha^−1^·10 years^−1^ (61), whereas results from our study found mortality of between 8-129 trees·ha^−1^ (Table 2), 2-44 times higher, occurring during just one extreme storm event. In addition, our estimates of mortality are conservative, since trees with minor damage or canopy damage may experience delayed mortality due to storm related injuries (4,39). In the Florida Panhandle alone, there have been 10 major hurricanes to make landfall since 1851 (67). Given the average return interval for a hurricane in the Florida Panhandle of 9-13 years (20), or 1 major hurricane every 2 years for the entire U.S. coastline (20), it is possible that historically hurricanes may have played a more important role in maintaining the population dynamics of longleaf pines than lightning at longer temporal scales.

### 4.3 Implications for management and restoration

For longleaf pine habitats affected by Hurricane Michael, active fire management will be critical to restoration (51–53,68). In all instances where trees were killed, by snapping or uprooting, the increased biomass on the ground contributes to fuels for fire and at a fine-scale change fire behavior by creating microsites that burn at hotter temperatures for longer amounts of time (69). In order to reintroduce fire to some of the more heavily damaged sites, low impact timber salvage will be necessary to remove dangerous fuel sources and open up the understory to promote fire contiguity while minimizing impact to the soil and understory (70,71). In sites where the mature trees are significantly reduced, such as Apalachee WMA, natural regeneration may no longer be possible and restoration should include planting of seedlings (22,60).

### 4.4. Conclusion

The current rate of loss of biodiversity and ecosystem services is unprecedented and is accelerating due to multiple interacting human stressors (72). In the NACP, storms of increasing strength and frequency pose a significant threat to the longleaf pine ecosystem and the numerous species that depend on it. Here we show that Hurricane Michael resulted in varying mortality on longleaf pines in the Florida Panhandle with the most severe impact resulting in catastrophic losses (92%) of mature canopy trees. This study focuses on the impact of Hurricane Michael in Florida, but the storm impacted most states within the NACP, all containing critical longleaf pine habitat. The increasing frequency of extreme stochastic events requires updating restoration and management plans for critical habitats (6). Managers and policy-makers attempting to mitigate climate change impacts need to account for potential unexpected losses and have contingency plans for responding to extreme disturbance events. Meeting current conservation targets will likely require protecting a larger extent of habitat than currently considered. The remaining extent of longleaf pine ecosystems exist in varying degrees of habitat integrity (16) and even protected high quality habitat is ecologically vulnerable to climate change. Moving forward, we must consider the implications of changing disturbance regimes due to anthropogenic climate change on the ecology of critical habitats.

## Acknowledgements

Thank you to the Florida Natural Areas Inventory and Florida Forest Service for data and access to the Longleaf Pine Ecosystem Geodatabase. We thank the various land agencies for continued management of the reference sites as well as access to perform our study, including the U.S. Fish & Wildlife Service, U.S. National Forest Service, and Florida Fish and Wildlife Conservation Commission. Thanks to Chad Anderson (FSU/FNAI) for use of his photo of Apalachee WMA. We thank those who assisted in collection of field data and in preliminary data exploration, including Gracie Rivera, Savana Roach, Ryan Slapikas and Shiqian (Kate) Wang.

